# Identification and characterization of a synaptic active zone assembly protein

**DOI:** 10.1101/2024.04.08.588536

**Authors:** J. Lützkendorf, T. Matkovic-Rachid, T. Götz, S. Liu, T. Ghelani, M. Maglione, M. Grieger, S. Putignano, L. Gao, N. Gimber, J. Schmoranzer, A. Stawrakakis, A.M. Walter, M. Heine, M.C. Wahl, T. Mielke, F. Liu, S.J. Sigrist

## Abstract

At presynaptic active zones (AZs), scaffold proteins play a crucial role in coordinating synaptic vesicle (SV) release and forming intricate nanoarchitectures essential for synaptic function. Despite their suspected importance, factors governing the assembly of nanoscale AZ scaffolds have remained elusive. Here, we identify “Blobby” as a novel regulator of AZ nanopatterning, localized within the AZ scaffold. Genetic loss of the extended Blobby protein led to aberrant accumulation of AZ scaffold proteins (“blobs”) and disrupted the nanoscale architecture of the AZ scaffold, resulting in a significant reduction in the packing density of voltage-gated Ca^2+^ channels at AZs, as observed through intravital single-molecule imaging. This disruption correlated with decreased evoked synaptic currents and SV release probability. Our findings suggest that Blobby plays a crucial role in switching the AZ scaffold into a state which allows to fine-tune the dynamic nanopatterning of Ca^2+^ channels to maintain proper release.

## Introduction

Active zones (AZs) are cellular platforms that govern presynaptic release function. These nanoscale-patterned macromolecular architectures serve as major signaling hubs for synaptic information transfer and storage ^1–4^. At AZ membranes, synaptic vesicles (SVs) are released and replenished at high speeds, with already subtle deficits in release and/or replenishment rates provoking a significant detrimental impact on the behavioral and thus organismal level. In ultrastructural and molecular terms, AZs are decorated by electron dense “scaffolds”, which form from extended cytoplasmic proteins belonging to only a few evolutionary conserved families. These scaffold structures are meant to help the recruitment of new SVs to AZ membrane ^5–7^ where they get biochemically primed for release at sites enriched for (m)unc13 family release factor proteins which operate in conjunction with nanoscopic clusters of conserved voltage-gated Ca^2+^ channels ^8–11^.

To execute their essential biological functions, AZ membranes must be meticulously patterned at the nanoscale level, typically within the range of tens of nanometers. This precise architecture is crucial for establishing both biochemical and topological connections between voltage-gated calcium channels and (m)unc13 clusters, fundamental for orchestrating synaptic vesicle release. Although the assembly of intricate protein structures often relies on the support of “assembly factors”, the identification of proteins that might specifically refine the assembly of AZ scaffolds and, by extension, the configuration of presynaptic release sites as well as the nanoscale distribution of voltage-gated Ca^2+^ channels has remained elusive. In other words, the precise mechanisms by which extended AZ proteins assemble into functional scaffolds to establish release sites and regulate synaptic vesicle release is still largely unknown. This said, genetic analyses in model organisms such as C. elegans, *Drosophila*, and mice have shed light on the critical role of the conserved ELKS family (BRP in *Drosophila*) in driving the AZ assembly process including the promotion of proper nanoscale clustering of voltage-gated Ca^2+^ channels ^12–20^.

In our study, we introduce a novel protein residing within the AZ scaffold of developing *Drosophila* synapses, termed “Blobby”. Comprising coiled coil and intrinsically disordered domains, Blobby lacks domains previously associated with synaptic function. Investigation at neuromuscular terminals during larval development revealed Blobby’s essential role in assembling the nanoscale protein architecture crucial for AZ release function, as evidenced by the near absence of proper electron dense structures covering the AZ membrane (“T-bars”). These structural deficits were accompanied by an abnormal nanoclustering of voltage-gated Ca^2+^ channels observed through intravital single-molecule imaging, obviously contributing to the compromised synaptic vesicle release. Our findings suggest that the intricate process of AZ assembly has driven the evolution of Blobby as a dedicated assembly-promoting protein.

## Results

### The novel protein Blobby accumulates in AZ scaffolds of developing *Drosophila* synapses

At *Drosophila* AZs, the ELKS-family scaffold protein BRP plays a crucial role in forming SV release sites by promoting the nanoscale localization of Unc13A, a member of the (m)unc13 family of proteins ^9, 10, 16^, and clustering Ca^2+^ channels at the AZ membrane via its N-terminal region ^16^. Additionally, BRP’s elongated shape, approximately 80 nm in length, facilitates SV targeting toward the SV release sites ^21–23^. BRP at *Drosophila* synapses fulfills the criteria of a “master AZ scaffold protein” whose availability determines the local, AZ and global, neuron-wide pools of Unc13A ^24^.

In our quest to identify additional proteins involved in the developmental assembly of AZ scaffolds, we conducted anti-BRP immunoprecipitations (Fig.1A) from *Drosophila* brain synaptosome preparations ^25^. As anticipated, we identified known BRP interactors, inter alia including RIM-BP ^15^ and Unc13A ^9^. However, the most prominent hit we detected was a previously uncharacterized protein encoded by the CG42795 locus. To capture an aspect of its mutant phenotype observed when the entire protein is eliminated described below, we henceforth refer to the encoded protein as “Blobby” (Fig. 1A).

**Figure 1.**
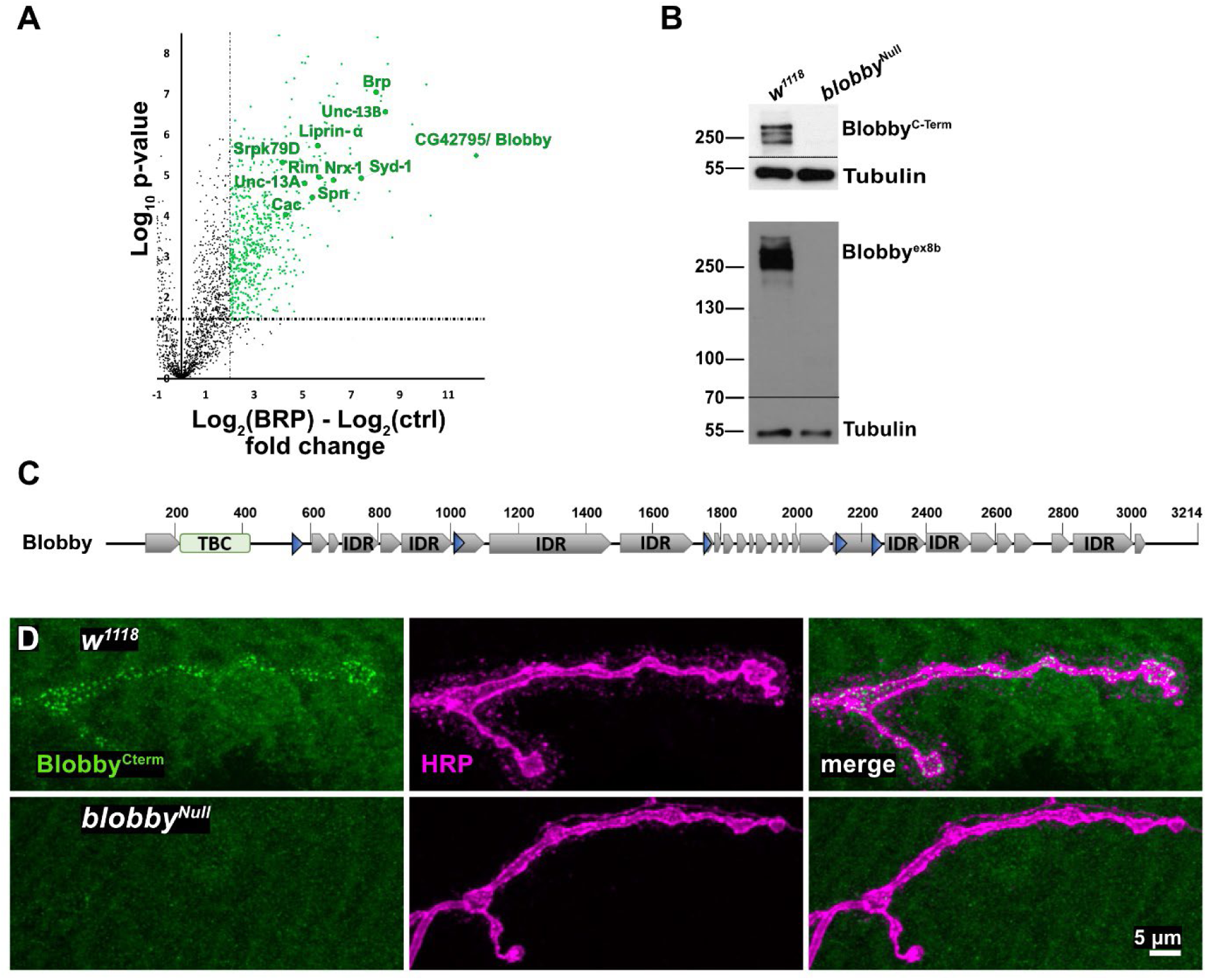
A novel extended protein enriched in AZ scaffold protein immunoprecipitates. **A)** Identification of CG42795/Blobby in BRP-immunoprecipitates. Volcano plot comparing BRP-immunoprecipitates with IgG control done in quadruplicate. X-axis: log2 fold-change value, Y-axis: – log10 of the p-value. **B)** Western Blot detecting Blobby reactivity in adult brain protein extracts of indicated genotypes, probed with both anti-Blobby^ex8b^ and anti-Blobby^C-term^. Anti-Tubulin probing serves as loading control. Prominent Blobby bands are detected in the wild type (*w^11^*^18^, first lane) but absent in *blobby*^Null^ mutants. **C)** Blobby domain structure depicting the predicted domains: TBC domain (yellow), coiled-coiled domains (blue), intrinsically disordered regions (IDR: grey)). **D)** Representative images of muscle 4 NMJs from third instar control and *blobby*^Null^ labeled with the Blobby^C-term^ antibody.

To detect the protein in brain homogenate Western blots (Fig. 1B), we generated antibodies against this novel factor. CG42795/Blobby encodes a sizable protein (several isoforms predicted by Flybase^26^ in the size range of 250-300 kD) distinguished by its extensive intrinsically disordered regions (IDRs) and coiled-coil (CC) domains (Fig.1C). Additionally, it contains a TBC putative Rab-GAP domain near its N-terminus, for which dendrogram analysis suggests a homology relation to the human protein TBC1D30 (Fig. S1A). Western probing in *w^11^*^18^ control animal brain extracts with either the ex8b Ab or the C-term Ab (for position of epitopes see Fig. S1B) detected several bands in the predicted size range of 250-300 KD (Fig. 1B).

To validate the specificity of our results, we generated *blobby*^Null^ mutants by simultaneously deleting two major segments of the *blobby* open reading frame via CRISP/Cas9 editing, as indicated in the gene locus map (Fig. S1A). The Western blot signals for both antibodies were eliminated in *blobby*^Null^ mutants (Fig. 1B).

Confocal microscopy analysis of larval neuromuscular junction (NMJ) terminals, identified by HRP staining, showed synaptic staining for Blobby, which was absent in *blobby*^Null^ mutant larvae, however (Fig. 1D).

Given the Co-IP of Blobby with BRP, and its immunofluorescence distribution, we wondered whether Blobby might be a new AZ scaffold protein. Indeed, confocal stainings of Blobby with either BRP or the AZ scaffold protein RIM-BP revealed a close co-localization between Blobby and two AZ scaffold components, BRP and RIM-BP (Fig.2A). Observing this obvious AZ scaffold localization, we sought to genetically confirm that presynaptic, motoneuron expressed Blobby is responsible for this AZ staining. To address this, we integrated a marker cassette flanked by KD recombinase sites into an exon of *blobby* locus using Crispr/Cas9 *(blobby-STOP-ALFA,* Fig. S1A). As expected, Blobby staining and ALFA-staining was abolished in this line (Fig.2B, upper row). However, after *ok6-Gal4* mediated KD-expression in motoneurons (and subsequent precise out-recombination of the KDRT-STOP-KDRT cassette), expression of ALFA-tagged Blobby was restored in ok6>blobby-ALFA (Fig. 2B). In the reverse experiment, the expression of postsynaptic (i.e. muscle) KD recombinase did not result in any Blobby staining (not shown). Thus, the Blobby signal at presynaptic AZs is obviously of motoneuron means presynaptic origin.

**Figure 2.**
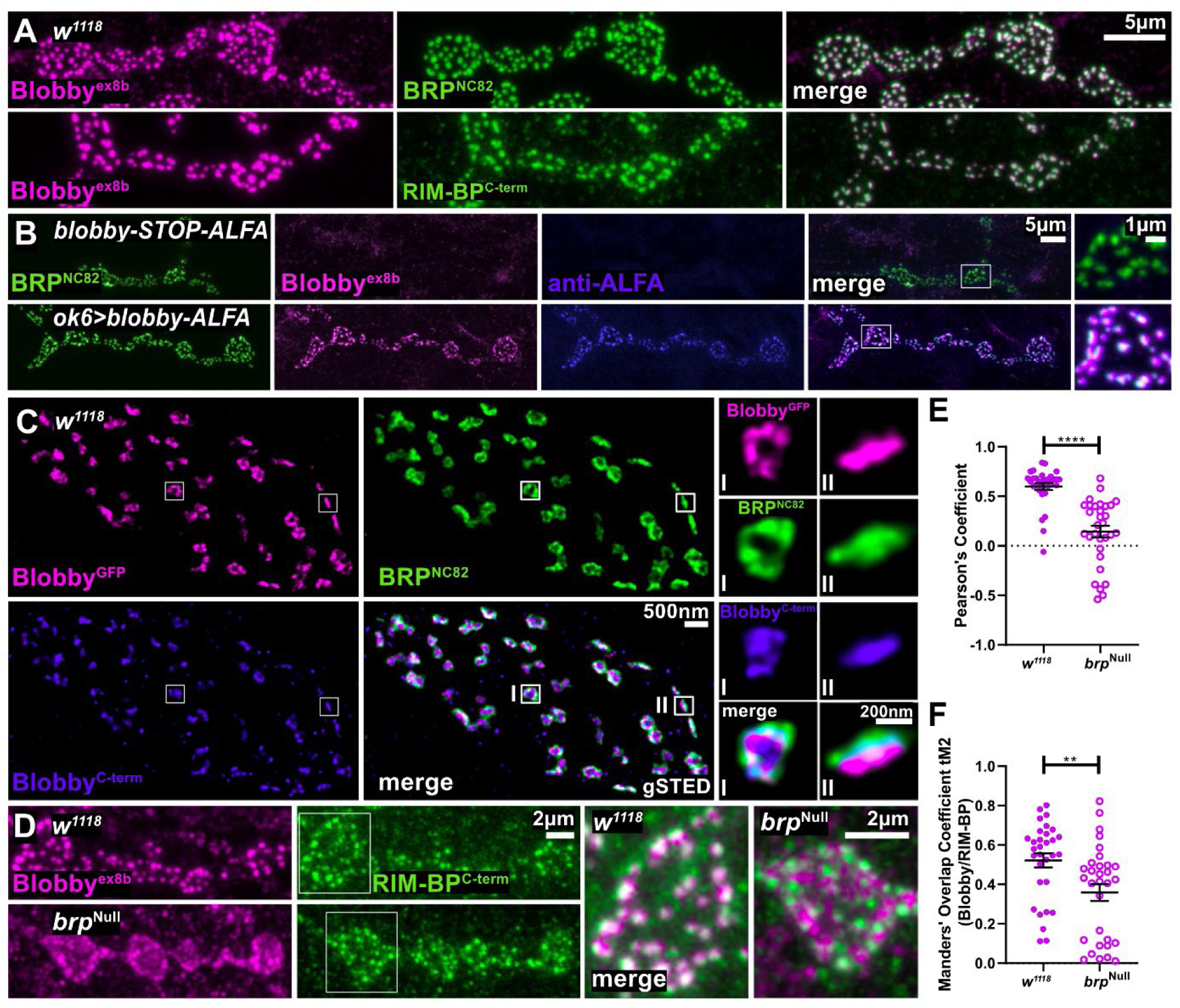
Blobby is tightly integrated into the AZ scaffolds of developing synapses. **A, B)** Shown are confocal images depicting muscle 4 neuromuscular junctions (NMJs) from third instar larvae of *w^1118^* labeled with the indicated antibodies. Further details on genotypes can be found in the main text. **C)** gSTED images of muscle 4 neuromuscular junctions of Blobby-GFP (magenta) animals stained for BRP^NC82^ (green) and Blobby^C-term^(blue). Magnified images of individual AZs imaged in planar (I) or vertical (II) orientation. Blobby^GFP^ - BRP^NC82^ 20 ± 2 nm, n=150 AZs, 4 animals; Blobby^GFP^ – Blobby^C-Term^ 14 ± 2 nm, n=47 AZs, 4 animals; Blobby^C-Term^ – BRP^NC82^ 17 ± 2 nm, n=47 AZs, 4 animals. **D)** Representative confocal images of muscle 4 NMJs from *w^111^*^8^ and *brp*^Null^ mutants stained with the Blobby^ex8b^ antibody (magenta) and RIM-BP^C-term^ (green), individual boxed boutons shown in higher magnifications. **E, F)** Quantification of overlap between BRP and Blobby^ex8b^ signals in controls and *brp*^Null^. **E)** Plot of Pearsońs coefficients: *w^111^*^8^ 0.59 ± 0.03, n=31, N=4; *brp*^Null^ 0.14 ± 0.05, n=31, N=4); **F)** Plot of Manders coefficients: *w^1118^* 0.5 ± 0.04, n=31, N=4; *brp*^Null^ 0.35 ± 0.04, n=31, N=4). Graphs show mean ± SEM. ****p<0.0001; **p<0.01 (Kolmogorov-Smirnov test was applied). N: number of animals, n: number of boutons.

To further explore the presynaptic AZ localization of Blobby in detail, we turned to time-gated stimulated emission depletion (gSTED) microscopy ^27^. We utilized the on-locus GFP-blobby line staining anti-GFP along with the Blobby^ex8b^ antibody and the BRP^NC82^ monoclonal antibody (Fig. 2C,D). This triple-gSTED experiment revealed that both Blobby epitopes were entirely confined within the BRP-positive AZ scaffold (Fig. 2C), particularly noticeable in planar, *en face* imaged AZs (see magnifications labeled with I). In vertically imaged AZs (labeled with II), average distances between all three epitopes were measured and found to be about 20 nm (for exact values, please refer to the figure legend) . Thus, Blobby protein is obviously integrated into the AZ-scaffold close to the BRP^NC82^ epitope. Given that the BRP^NC82^ epitope marks the distal end of the T-Bar AZ scaffold in a distance of about 70 nanometers from the AZ localized SV release sites ^28^, Blobby is obviously a novel AZ scaffold component tightly co-localizing with BRP in the more distal AZ scaffold.

### BRP is crucial for the AZ scaffold incorporation of Blobby

We next investigated whether the localization of Blobby within AZ scaffolds might rely on BRP. Thus, we compared control with *brp*^Null^ mutant NMJ terminals by performing co-immunofluorescence stainings of Blobby^ex8b^ with RIM-BP^C-term^ (Fig. 2D-F). RIM-BP staining was selected because it can detect AZ scaffolds even in the absence of BRP ^15^. Confocal images of controls as expected (Fig. 2A) showed that Blobby co-localized with RIM-BP (Fig. 2D). However, the degree of co-localization appeared clearly reduced at the *brp*^Null^ mutant terminals (Fig. 2E merged magnifications to the right), evident also after quantification using two distinct co-localization scores (Fig. 2E,F). Thus, Blobby’s localization within AZ scaffolds appears to be impaired in the absence of BRP. Notably, however, in the absence of BRP the bulk of Blobby still reached a position close to the AZ scaffolds implying that its transport is largely independent of BRP (also see discussion).

### Reduced AZ number and ectopic aggregation of scaffold proteins at *blobby* terminals

Subsequently, we proceeded to investigate a putative role of Blobby in NMJ development and presynaptic AZ assembly. Thus, we stained larvae for BRP and the postsynaptic glutamate receptor GluRIID (Fig. 3A) ^29^. NMJ area (Fig. 3B) and AZ number (Fig. 3 C; via BRP spot numbers) were both significantly reduced at *blobby*^Null^ larval NMJs. The AZ area density was not significantly changed (Fig. 3D). Upon closer inspection of the BRP signals, we observed unusually large BRP aggregates, which we termed “blobs” (Fig. 3A, arrow). This was the first phenotype we observed and thus motivated the choice of the name *blobby*. Unlike the AZ BRP signals, these blobs were not positioned in apposition to postsynaptic glutamate receptors, as evident from the lack of overlay between BRP and GluRIID staining in confocal image projections (Fig. 3A, arrow). Quantification based on the lack of BRP/GluRIID overlay indeed uncovered a significant accumulation of this obviously ectopic material in *blobby*^Null^ mutants (Fig. 3E).

**Figure 3.**
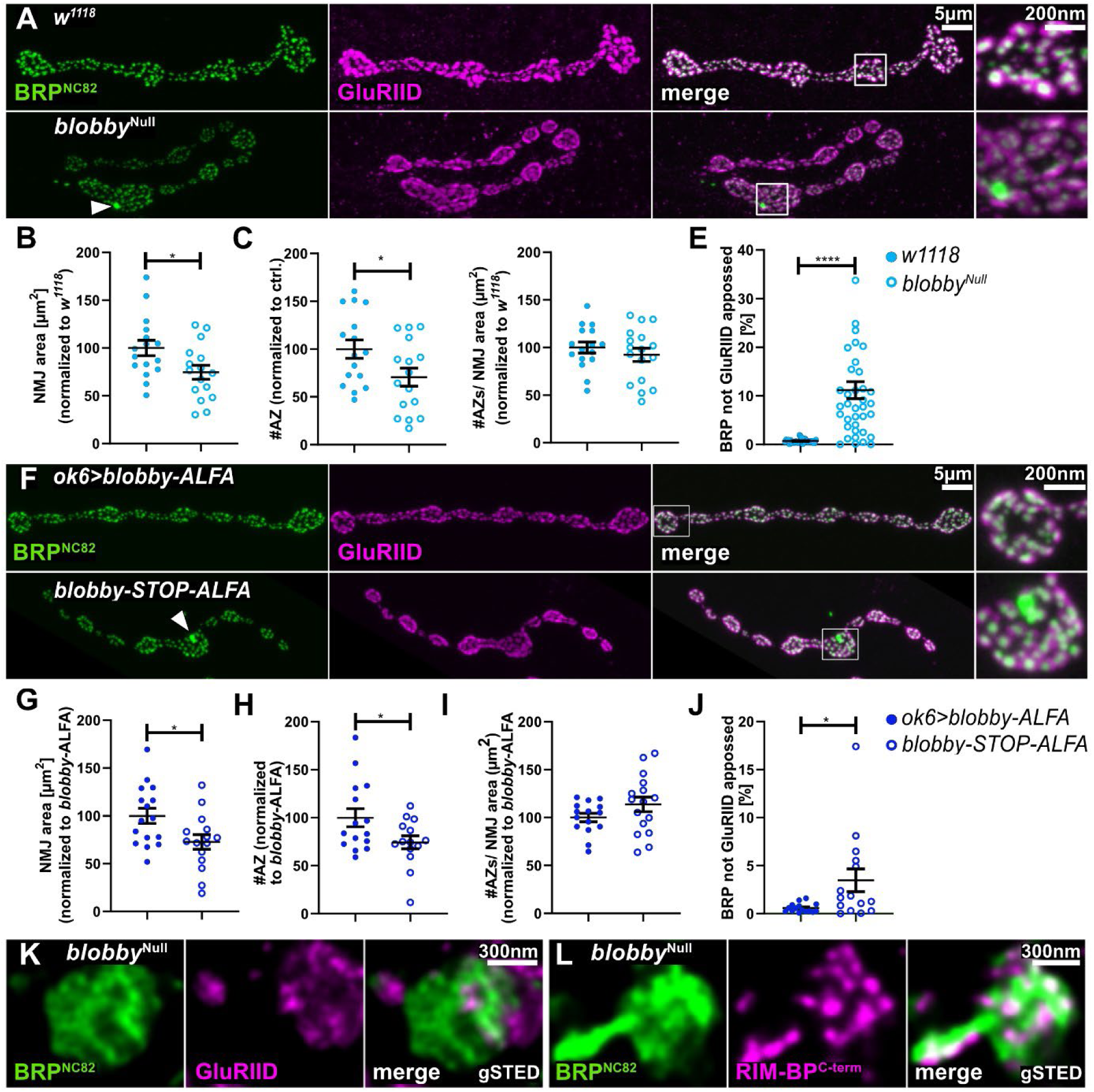
Analysis of *blobby* mutant NMJs via confocal microscopy. **A-J**: Confocal imaging analysis comparing larval NMJs of *blobby*^Null^ animals to controls (A-G) and *blobby-*STOP*-*ALFA to *ok6>blobby-*ALFA (F-J). **A, F)** Representative confocal images of muscle 4 NMJ images of third instar larvae stained with BRP^NC82^ (green) and GluRIID (magenta) of indicated genotypes; **B-E) and G-J)**: Quantification of NMJ morphology and AZ numbers of indicated genotypes. **B, G)** Normalized NMJ area (in µm^2^): *w^1118^* 100 ± 8.1, n=16, N=4; *blobby*^Null^ 74.58 ± 7.26, n=16, N=4; *ok6>blobby-*ALFA 100 ± 7.9, n=16, N=4; *blobby-*STOP*-*ALFA 72.78 ± 7.66, n=15, N=4; unpaired student t-test); **C, H)** Number of AZs identified as discrete BRP positive clusters per NMJ, values normalized to wild type: *w^1118^* 100 % ± 9,6 % n=16, N=4; *blobby*^Null^: 70.7 % ± 9.4 %, n=16, N=4; *ok6>blobby-*ALFA: 100 % ± 9.49 %, n=15, N=4; *blobby-*STOP*-*ALFA 74.29 % ± 6.78 %, n=14, N=4, unpaired student t-test. **D, I)** Number of AZs normalized to NMJ area from projected images, values normalized to respective controls: *w^1118^* 100 ± 5.71, n=16, N=4; *blobby*^Null^ 92.35 ± 6.89, n=17, N=4; *ok6>blobby-*ALFA 100 ± 4.35, n=15, N=4; *blobby-*STOP*-*ALFA 113.7 ± 7.7, n=16, N=4, unpaired student t-test) **E, J)** Quantification of ectopically distributed BRP signal as BRP signal not apposed to GluRIID signals in projected confocal images; *blobby*^Null^ in comparison to control: *w1^118^* 0.7% ± 0.07%, n=32, N=5; *blobby*^Null^ 11.2% ± 1.7%, n=38, N=5); *ok6>blobby-*ALFA 0.59% ± 0.1%, n=15, N=4; *blobby-*STOP*-*ALFA 3.5% ± 1.2%, n=15, N=4, Kolmogorov-Smirnov test). Graphs show mean ± SEM. ****p<0.0001; *p<0.05. N: number of animals, n: number of NMJs analyzed. **K-L)** Representative time-gated (g)STED images highlighting BRP^NC82^-positive aggregates (“blobs”) of *blobby*^Null^ mutants in synaptic boutons stained with **(K)** BRP^NC82^ (green) showing apposed signal of postsynaptic GluRIID (magenta) and **(L)** BRP^NC82^ (green) and RIM-BP^C-term^ (magenta) indicating co-labeling of both epitopes in the blobs.

We then in identical experiments tested the animals without *(blobby-STOP-ALFA)* and with (*ok6>blobby-ALFA*) motoneuron-specific expression of Blobby (Fig. 3F-J). Significant reductions of NMJ area (Fig. 3G) and AZ number (Fig. 3H) were also found in *blobby-STOP-ALFA* relative to *ok6>blobby-ALFA.* Moreover, blobs were specifically observed in *blobby-STOP-ALFA* but not in *ok6>blobby-ALFA* animals with presynaptic Blobby expression (for quantification see Fig. 3J).

We conducted a final examination of the ectopic BRP blobs using gSTED microscopy (Fig. 3K,L). As anticipated, these aggregates were not found to be associated with postsynaptic GluRIID signals in 2-D projected images (Fig. 3K). However, RIM-BP, “stochiometric” component of *Drosophila* AZ scaffolds^15^, was present within these blobs with a similar topology as in mature AZs (Fig. 3L). These data suggest that Blobby is expressed and functionally relevant within the presynaptic neuron to ensure a robust targeting of AZ scaffold components into AZs, and to prevent the accumulation of ectopically assembled AZ scaffold material (also see discussion).

### Decreased transmission and reduced SV release probability at Blobby lacking synapses

To evaluate Blobby’s impact on synaptic vesicle (SV) release on larval NMJ synapses, we conducted two-electrode voltage clamp recordings to measure both action potential-evoked and spontaneous miniature SV release. We again compared the two genetic scenarios involving the loss of presynaptic Blobby: *blobby*^Null^ versus isogenic control (Fig. 4A-H) and also *blobby-STOP-ALFA* versus *ok6>blobby-ALFA* (Fig. 4I-P). Action potential evoked postsynaptic excitatory junctional currents (eEJCs) were significantly reduced at NMJs lacking Blobby (Fig. 4A, B, I, J). Quantal contents, indicative of the number of SVs released per action potential, were then calculated by dividing eEJC values by the respective miniature excitatory junctional amplitudes (mEJCs; Fig. 4G, O). Both Blobby-deficient genetic backgrounds showed strongly and significantly reduced quantal contents (Fig. 4C, K), demonstrating a decrease in the efficacy of evoked SV release in the absence of Blobby.

**Figure 4.**
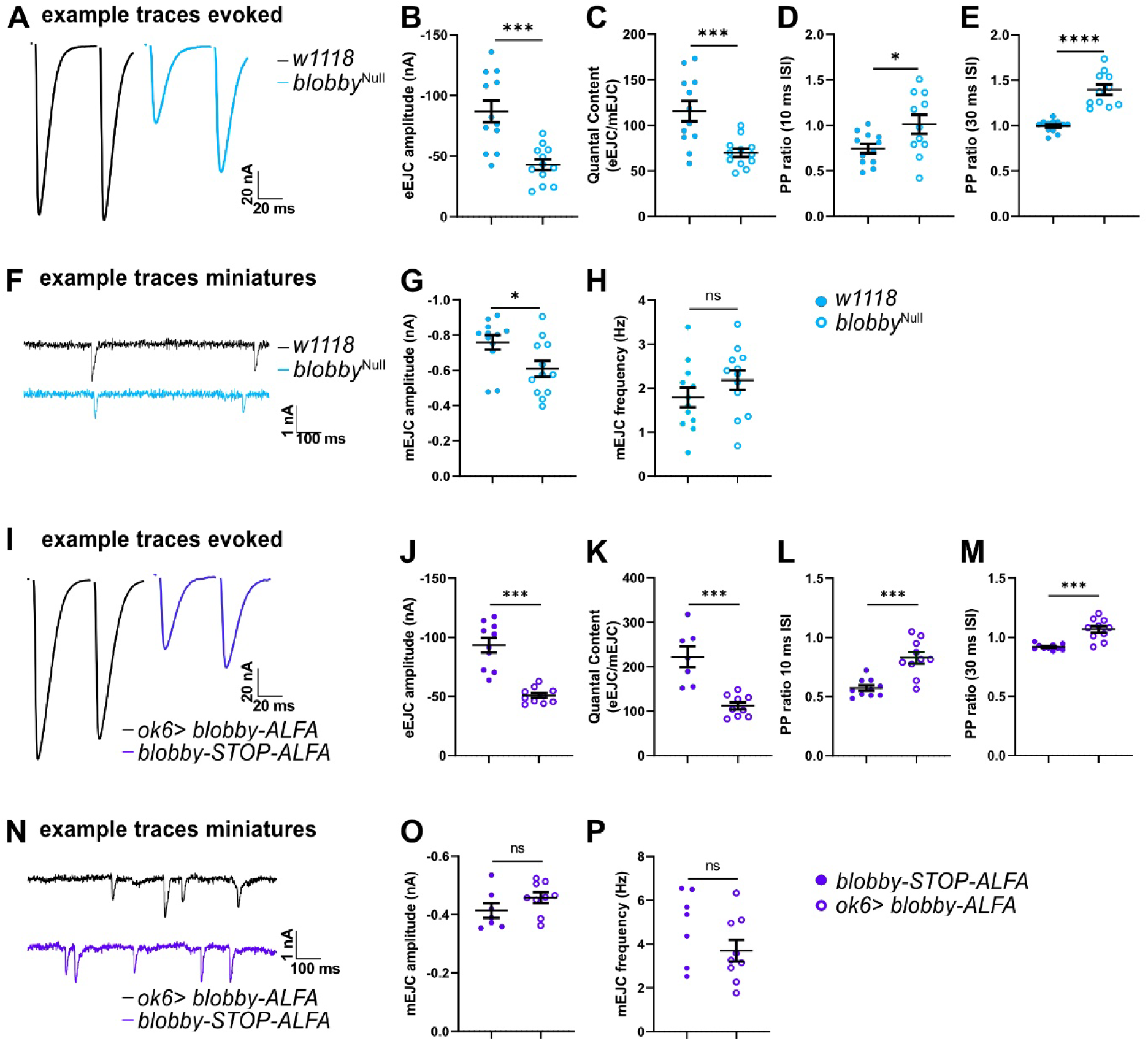
Electrophysiological analysis of *blobby* mutant NMJs. **A-P)** Two-electrode voltage clamp electrophysiological recordings comparing third instar larvae NMJs of *blobby*^Null^ animals to controls (A-H) and *blobby-*STOP*-*ALFA to *ok6>blobby-*ALFA (I-P). **A, I)** Example traces of evoked responses at interpulse interval of 30 ms. **B, J)** eEJC amplitudes (*w^1118^* -86.91 ± 8.96 nA, n=12; *blobby*^Null^ -43.03 ± 4.40 nA, n=12; ok6>*blobby-*ALFA -93.42 ± 6.13 nA, n=10; *blobby*-STOP-ALFA -45.73 ± 3.40 nA, n=10). **C, K**) Quantal contents (*w^1118^* 115.70 ± 11.09, n=12; *blobby*^Null^ 69.88 ± 4.49, n=12; ok6>*blobby-*ALFA 222.20 ± 23.23, n=7; *blobby*-STOP-ALFA 87.74 ± 6.96, n=10). **D, L**) Paired-pulse ratio at interpulse intervals of 10 ms. (*w^1118^* 0.74 ± 0.05, n=12; *blobby*^Null^ 1.01 ± 0.10, n=11; ok6>*blobby-*ALFA 0.57 ± 0.02, n=10; *blobby*-STOP-ALFA 1.03 ± 0.05, n=10). **E, M**) Paired-pulse ratio at interpulse intervals of 30 ms. (*w^1118^* 0.99 ± 0.02 n=12; *blobby*^Null^ 1.39 ± 0.06, n=11; ok6>*blobby-*ALFA 0.92 ± 0.01, n=10; *blobby*-STOP-ALFA 1.30 ± 0.02, n=10). **F, N)** Example traces of miniature responses. **G,O)** mEJC amplitudes (*w^1118^* -0.75 ± 0.04 nA, n=12; *blobby*^Null^ -0.60 ± 0.05 nA, n=12; ok6>*blobby-*ALFA -0.41 ± 0.03 nA, n=7; *blobby*-STOP-ALFA -0.55 ± 0.05 nA, n=10). **H, P)** mEJC frequencies (*w^1118^* 1.79 ± 0.22, n=12; *blobby*^Null^ 2.18 ± 0.22, n=12; ok6>*blobby-*ALFA 4.83 ± 0.62, n=7; *blobby*-STOP-ALFA 1.66 ± 0.31, n=10). Graphs show mean ± SEM. An unpaired t-test was applied, *p < 0.05; ***p<0.001; ****p < 0.0001 ns = not significant. n represents a single cell. Four to six animals are analyzed with one or two cells/animal.

One potential explanation for this decrease in SV release could be an impairment of SV release probability resulting from the loss of Blobby function. Paired-pulse facilitation or depression, in response to rapid presynaptic stimulation, serves as a means to estimate SV release probability, with increased facilitation indicating decreased release probability. In both *blobby*^Null^ compared to the isogenic control and *blobby-STOP-ALFA* versus *ok6>blobby-ALFA*, paired pulse ratios were significantly elevated across different interpulse intervals (Fig. D, E, L, M). To understand the underlying mechanisms behind the reduced SV release probability and SV release, we conducted further analysis on the nano-organization of individual AZs in *blobby*^Null^.

### Lack of Blobby disrupts the nanoscale organization of AZs

At the confocal level, we observed a reduction in the levels of BRP per AZ at *blobby*^Null^ NMJs in two distinct quantifications (Fig. S2 A,B). We then went on with gSTED analysis. Typically, under gSTED, the labeling of the BRP-C-term with the BRP^NC82^ monoclonal antibody reveals “smooth, ring-like structures” at planar imaged AZs (Fig. 2C magnification in I), as extensively described in previous studies ^30^. However, at *blobby*^Null^ NMJs, we observed a notably atypical pattern in the nanoscale spacing and distribution of BRP (Fig. 5A). Frequently, triangular and “zig-zag-like” arrays were observed (Fig. 5A, arrows), which were not observed in controls. A perimeter measurement, which distinguishes such zig-zag patterns from the typical round appearance, was significantly elevated in *blobby*^Null^ (Fig. 4B). We then gSTED co-imaged RIM-BP (Fig. 4C) and Unc13A (Fig. 4D) with BRP. While both proteins were still present at the defective BRP AZ scaffolds of the *blobby* mutant, their localization pattern followed the disturbed BRP nanopattern (Fig. 4C,D).

**Figure 5.**
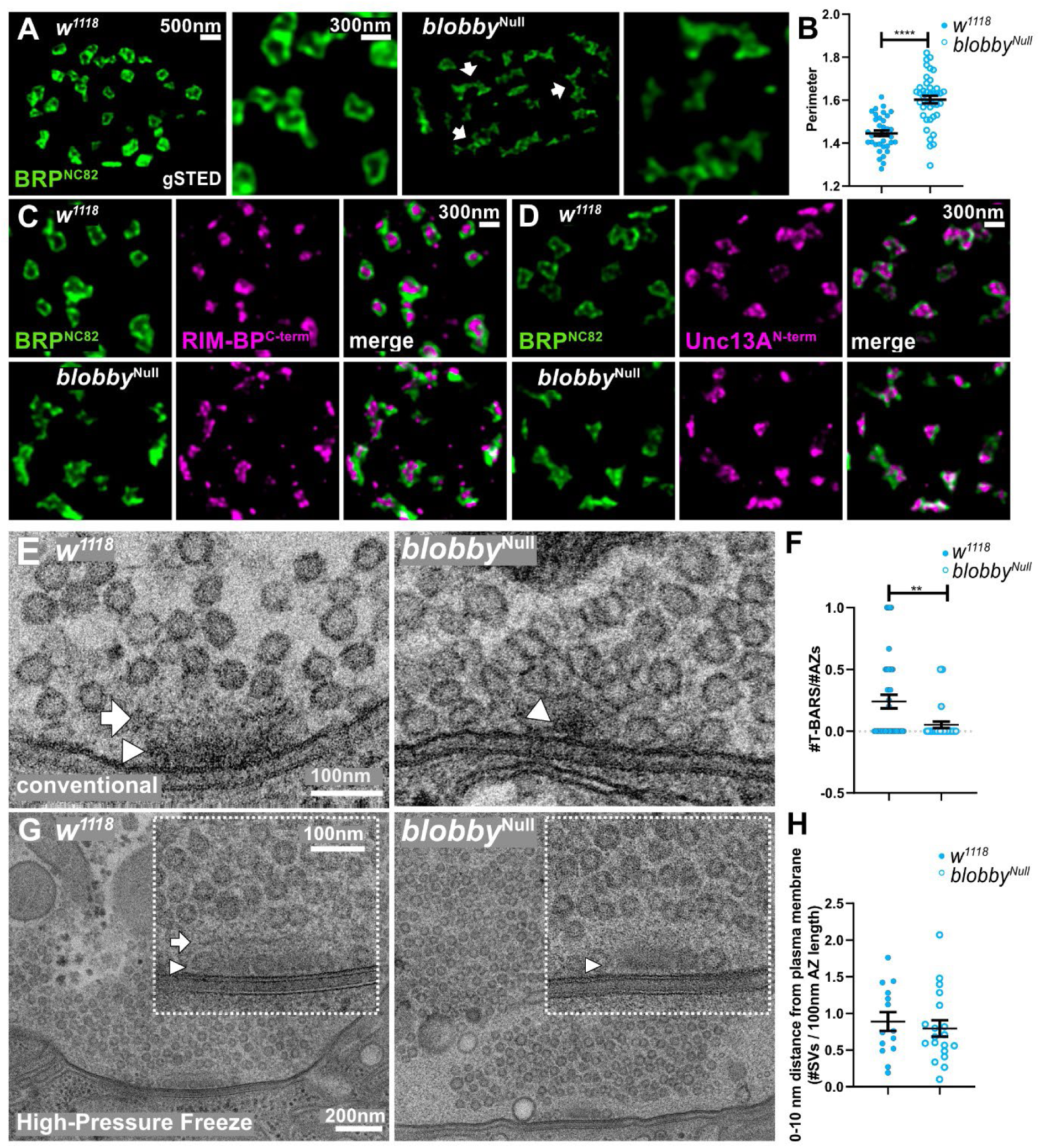
STED and EM characterization of *blobby*^Null^ defective AZ scaffolds. **A)** gSTED images of muscle 4 neuromuscular junctions of *w^1118^* and *blobby*^Null^ AZs stained for BRP. Arrows indicate disrupted “zig-zag-like” BRP scaffolds of *blobby*^Null^ mutants. Scale Bar of overview bouton: 500 nm. Scale Bar of magnified images of individual AZs: 300nm. **B)** Quantification of third instar larvae AZ nanoarchitecture. The perimeter of acquired gSTED images from *w^1118^* and *blobby*^Null^ were analyzed (*w^1118^* 1.40 ± 0.01, n=37; *blobby*^Null^ 1.60 ± 0.02, n=42. Graph shows mean ± SEM. Data distribution is normal according to the D’Agostino and Pearson omnibus normality test. One-way ANOVA was applied, using Tukey post-test, ***, p < 0.001; ****p < 0.0001. N: number single boutons analyzed in gSTED. **C, D)** gSTED images of muscle 4 neuromuscular junctions of *w^1118^* and *blobby*^Null^ AZs co-stained for BRP (green) and **C)** RIM-BP (magenta) and **D)** Unc13A (magenta). Scale Bar: 300 nm. **E, G)** Representative electron microscopy (EM) images of *w^1118^* and *blobby*^Null^ mutant boutons from **E)** conventional standard transmission EM using glutaraldehyde fixation and **G)** high-pressure freeze EM. Arrow labels T-Bar roof, arrowhead T-Bar pedestal. Scale Bar for overview is 200 nm, for zoom 100 nm. **F)** Quantification of morphologically identified T-Bar numbers per AZ section (*w^1118^* 0.24 ± 0.05, n=38, N=80; *blobby*^Null^ 0.05 ± 0.02, n=33; N=76). Graph shows mean ± SEM. N: number of boutons; N: number of AZs. 3 animals per genotype were analyzed. A Kolmogorov-Smirnov test was applied. **H)** Quantification of synaptic vesicle densities in a distance from 0 -10 nm from the AZ plasma membrane per 100 nm AZ (*w^1118^* 0.88 ± 0.12, n=14; *blobby*^Null^ 0.79 ± 0.10, n=19). Graph shows the mean ± SEM. N: number of boutons from 3 animals per analyzed genotype. A Kolmogorov-Smirnov test was applied.

We then subjected *blobby*^Null^ NMJ terminals to electron microscopy (EM). In standard transmission EM, when glutaraldehyde fixation is employed, *Drosophila* presynaptic AZ membranes typically appear covered by electron-dense structures known as T-bars, representing a structural imprint of the protein-rich AZ scaffold ^31^, as can be seen in Fig. 5E (arrow labels T-bar roof, arrowhead T-bar pedestal). Irregularities in the nanoarchitecture of BRP would thus likely impact the organization of T-bars. However, analysis of *blobby* mutants revealed a near absence of properly shaped T-bars, with irregularly shaped electron-dense structures frequently observed instead (Fig. 5E, arrowhead).

High-pressure freeze electron microscopy (HPF-EM) analysis is utilized to preserve the physiological state of biological samples by rapidly freezing them under high pressure^9, 32–34^. This technique ensures minimal damage to cellular structures and provides detailed insights into the ultrastructure of specimens. Also under HPF-EM, T-bar structures could be readily identified at control AZs (Fig. 5G left, pedestal marked by an arrowhead and roof structure marked by an arrow) but not at *blobby*^Null^ AZs (Fig. 5G right, residual structure marked by arrowhead).

Hence, Blobby plays a pivotal role in facilitating the precise nanoscale organization of the BRP scaffold, as supported by evidence from both STED microscopy and electron microscopy.

### *In vivo* single particle imaging: defective AZ Ca^2+^ channel clustering at *blobby* synapses

HPF-EM analysis also facilitates the quantification of physiological synaptic vesicle (SV) numbers and their distributions relative to the AZ membrane ^35^. We aimed to investigate whether SV release probability correlated with a decreased number of SVs in close proximity to the AZ plasma membrane. However, counting analysis revealed only slight and non-significant reductions in SV numbers within 10 nm of the AZ membrane, both within 0-5 nm (Fig. S2 C) and 0-10 nm (Fig. 5H) bins. Therefore, deficits in SV recruitment, biochemical priming, and docking may not solely account for the severe deficits in SV release observed in the absence of Blobby AZ function. These functional deficits may instead be attributed to alterations in the precise nanopatterning of voltage-gated Ca^2+^ channels.

The Ca^2+^ channel α-subunit Cacophony (Cac), evolutionary homologous to the Ca_v_2.2. family, governs SV release at *Drosophila* AZs ^36^, and is regulated by BRP ^30^. Despite their low levels at *Drosophila* AZs, recent advancements in live single-molecule sptPALM imaging of fully functional mEOS-labeled Cac^37^ offer high contrast, enabling precise localization of individual Cac Ca^2+^ channel molecules at intravitally imaged larval NMJs ^16^.

Utilizing Dual-Delaunay Triangulation Tessellation, a segmentation method based on cumulative localizations ^16, 38^, Cac sptPALM data retrieved from NMJ AZs revealed robust identification of a central “nanocluster” (NC), surrounded by an area of significant Cac labeling density termed the AZ (Fig. 6A) ^16^.

**Figure 6.**
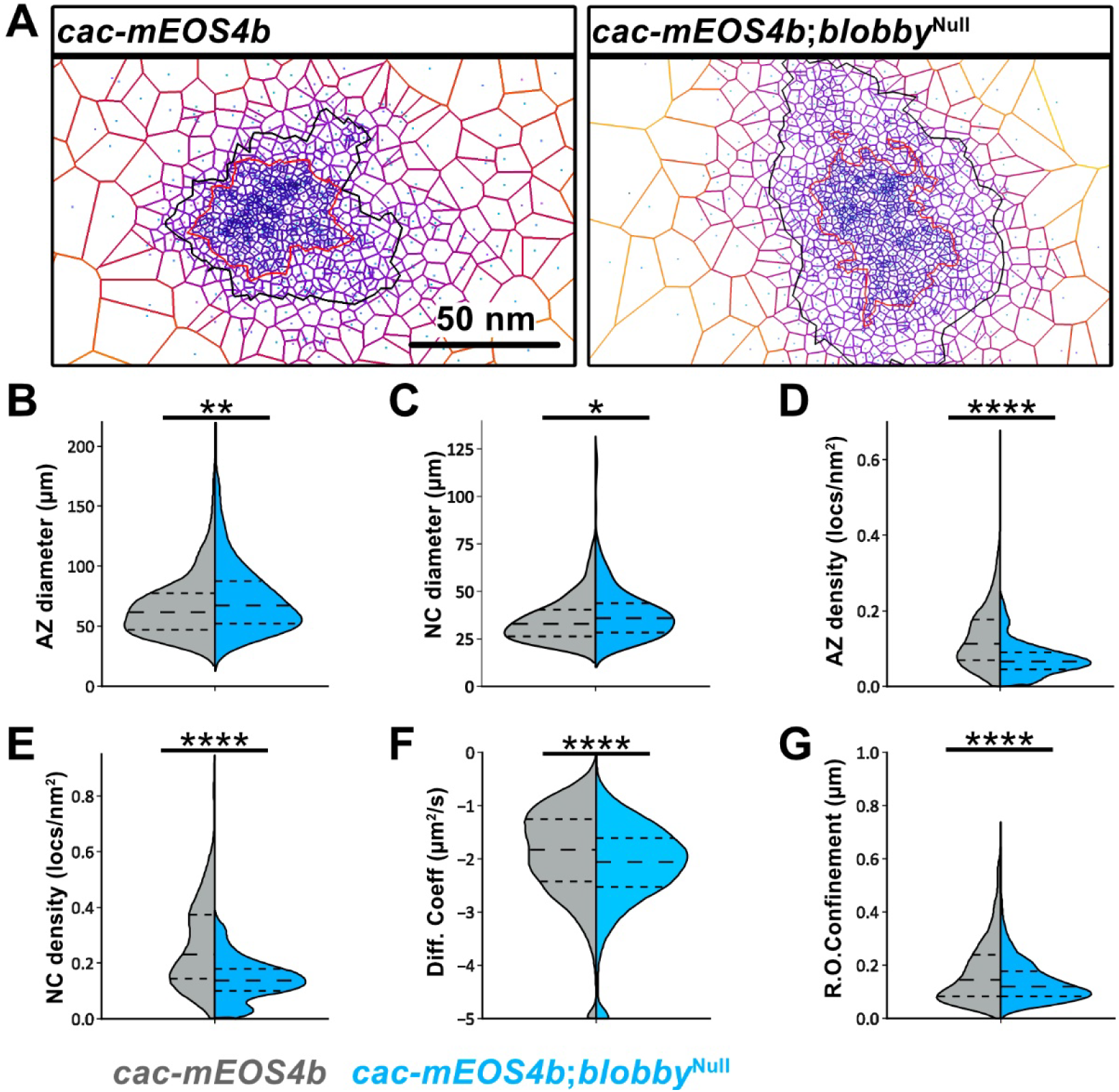
Intravital single molecule imaging of Cacophony at *blobby* mutants AZs. **A-E)** Live sptPALM imaging of *cac-mEOS4b* (control) and *cac-mEOS4b*; *blobby*^Null^ mutant were imaged in 1.5mM Ca^2+^extracellular imaging buffer. For details see^16^. **A, B)** Representative boutons displaying the live cumulative distribution of Cac localization as a tessellation map of an individual synapse. Scale Bar 50nm. **B)** Tessellation analysis of AZ diameter, **C)** nano-cluster (NC) diameter, **D)** Cac localization density in AZs and **E)** NCs. **F)** Live diffusion coefficient and **G)** radii of confinement derived from the mean square displacement of single live imaged Cac molecules in *cac-mEOS4b* (grey) and *cac-mEOS4b*; *blobby*^Null^ (blue). Data were pooled from 8-10 NMJs from 3-5 animals for each condition from which 242 AZs (*cac-mEOS4b* control), 254 AZs (*cac-mEOS4b*;*blobby*^Null^). Individual *Blobby*^Null^ experiments were conducted with concurrent controls. PALM Data distribution was statistically tested with Kolmogorov-Smirnov test. Statistical significance is denoted as asterisks: *p < 0.05; **p < 0.01; ***p < 0.001; ****p < 0.0001.

Using sptPALM imaging, we analyzed Cac localization at *blobby*^Null^ NMJs. Trajectories of single Cac molecules were recorded, and cumulative localization distributions were analyzed via tessellation (Fig. 6A). Although the total number of Cac localization events per AZ remained unchanged, we observed an expansion of both the AZ (black line) and NC (red line) areas, indicating a “looser packing” of Cac channels. Quantitative analysis revealed an increase in AZ and NC diameter (Fig. 6B,C), coupled with reduced Cac channel localization densities in both areas (Fig. 6D,E). These results suggest a deficiency in Cac Ca^2+^ channel organization at the single-molecule and nanopatterning level, likely contributing directly to the observed decrease in SV release probability.

We quantified Cac Ca^2+^ channel single molecule motility by analyzing diffusion coefficients (Fig. 6F) and radii of confinement from individual trajectories (Fig. 6G). Both parameters showed a significant *reduction* at Blobby-deficient AZs. In contrast, we previously found that AZs lacking BRP exhibit a significant *increase* in both parameters ^16^. Hence, the deficit in voltage-gated Ca^2+^ channel clustering observed in Blobby-deficient AZs is not solely attributable to reduced BRP levels within the AZ. Instead, we propose that Blobby is essential for transitioning the AZ scaffold into a functional state crucial for dynamically stabilizing the proper nanoscale arrangement of Ca^2+^ channels, thereby ensuring effective SV release.

## Discussion

Here, we present the discovery and characterization of Blobby, a novel protein found within AZ scaffolds at developing *Drosophila* synapses. Genetic ablation of Blobby disrupted the nanoscale molecular organization of AZs, as evidenced by i) abnormal distribution of the BRP AZ scaffold protein (Fig. 5A), ii) altered electron-microscopic appearance of the AZ scaffold (“T-bar”) (Fig. 5E,F), and iii) reduced density of voltage-gated Ca^2+^ channels at the AZ membrane, observed through sptPALM live intravital tracking (Fig. 6). Additionally, the absence of Blobby frequently led to an aberrant accumulation of ectopic AZ scaffold material (“blobs”) within presynaptic boutons.

Action potential-triggered synaptic vesicle (SV) release was notably diminished in the absence of Blobby, likely partially due to the reduced number AZs in this mutant. However, the elevated paired-pulse ratios suggest a decrease in SV release probability at Blobby-deficient AZs, aligning with observed defects AZ nanopatterning of voltage-gated Ca^2+^ channels. Thus, while the absence of AZ scaffold protein within ectopic blobs may contribute to fewer AZ formations, Blobby’s assembly function appears crucial for proper AZ scaffold nanopatterning, consistent with its direct localization within developing AZ scaffolds. Indeed, these effects on release probability cannot solely be attributed to the loss of BRP scaffold material. This is supported by the analysis of NMJs lacking the small GTPase Arl8, which exhibit reduced BRP levels and AZ numbers yet maintain unchanged paired pulse ratios, suggesting unaffected SV release probability^39^. Hence, our findings highlight the crucial involvement of Blobby in refining the intricate texture of AZ scaffold assembly during the development of synapses, vital for transitioning the AZ scaffold into a functional state necessary for precise nanoscale assembly. This process ensures the accurate arrangement of voltage-gated Ca^2+^ channels at the nanoscale, thereby facilitating effective SV release.

The mechanism(s) by which Blobby orchestrates the effective assembly and proper nanopatterning of AZs is an intriguing question. Its incorporation into the scaffold relies on BRP, and Blobby is obviously abundant in anti-BRP immunoprecipitates (Fig. 1A). The BRP/ELKS AZ scaffold family proteins, crucial for both the developmental assembly, functional operation but also plasticity of presynaptic release sites, contain extended CC domains and intrinsically disordered regions (IDRs) ^40, 41^. Blobby similarly is dominated by predicted IDRs and several CC domains (Fig. 1C). These features in all likelihood play a central role in the intricate process of AZ assembly, with IDRs promoting Liquid-Liquid Phase Separation (LLPS), which is obviously critical for the ordered and efficient assembly of the scaffold ^3, 42–46^. Recent findings from C. elegans suggest that a developmental liquid phase formed by the BRP homologue ELKS together with Syd-2/Liprin-α is essential for the initial assembly of the synaptic AZ scaffold before maturation into a stable structure ^47^. It is tempting to speculate that at nascent AZs in *Drosophila*, the IDRs of Blobby might play a role in maintaining BRP/ELKS-containing condensates in a liquid state during a crucial step of the AZ assembly process. An inability to sustain BRP in a liquid/soluble state is not unlikely to account for the formation of BRP blobs in the absence of Blobby. Intravital imaging studies have shed light on the dynamics of BRP accumulation at individual assembling *Drosophila* NMJ AZs, revealing a process that gradually unfolds over many hours ^28, 48^. Indeed, Blobby accumulation may be required throughout this developmental period to orchestrate the coordinated transfer of scaffold material into the growing AZ structures.

Phosphorylation has emerged as a key regulatory mechanism in controlling AZ assembly, likely also by modulating Liquid-Liquid Phase Separation (LLPS) formation. This post-translational modification is thought to dynamically regulate the interactions between scaffold proteins, thereby influencing their ability to undergo phase separation and assemble into functional AZ scaffolds. However, the specific phosphorylation events and their downstream effects on AZ assembly and function remain an active area of investigation ^42, 49–52^. A specific SRPK family protein kinase, SRPK79D, has been identified as a key player in the efficient transport and assembly of BRP^53, 54^. Notably, a homologous SRPK in rodent hippocampal neurons has been implicated in both AZ assembly and functional maturation ^51, 55^. In *Drosophila*, SRPK79D promotes the N-terminal phosphorylation of a region within the BRP IDR, crucial for efficient AZ incorporation^52^. This phosphorylation event prevents premature deposition of BRP, akin to observations in *blobby* mutants. The aberrant accumulations also contain RIM-BP, similar to the blobs in *blobby* mutants we observe in this study. However, only the BRP 190 KD isoform with the N-terminal IDR subject to SRPK phosphorylation is essential for blob formation in *srpk*79D^VN^ mutants. Hence, BRP acts as a “driver” protein for scaffold assembly, with RIM-BP and Unc13A as client proteins. This again suggests that maintaining BRP in a soluble “liquid” state might be crucial to prevent premature aggregate/blob formation and ensure coordinated AZ incorporation, and Blobby is now an obvious candidate to mediate this.

The identification of Blobby signifies the emergence of a specialized assembly factor, suggesting the development of regulatory mechanisms tailored to the demands of the AZ assembly process. An in-depth biophysical analysis of this interaction is warranted to further elucidate the underlying mechanisms governing AZ assembly. Such investigations would provide valuable insights into the molecular dynamics and structural features driving the formation and stabilization of AZ scaffolds, ultimately enhancing our understanding of synaptic development, plasticity and function.

## Acknowledgments

We acknowledge the assistance of the core facility BioSupramol supported by the DFG and the research center SupraFAB. We thank Ryota Fukaya for comments; Berit Söhl-Kielczynski and Christian Rosenmund for providing us access to the HPF machine and for support in performing the HPF experiments; Heike Stephanowitz for excellent technical assistance in performing mass-spectrometry experiments.

## Funding

This work was supported by grants from the Deutsche Forschungsgemeinschaft (DFG: German research Foundation):

Sonderforschungsbereich (SFB) 958

Neuronex Program IRG1

## Materials and Methods

### Fly husbandry, stocks and handling

Fly strains were reared under standard laboratory conditions and raised at 25°C on semi-defined conventional cornmeal-agar medium (Bloomington recipe) with 60-70 % humidity at 25°C. For electrophysiological experiments only male larvae were used. For all other experiments, both male and female third instar larvae or adult flies were used. For proteomics/WB, confocal and STED stainings, electrophysiology the following genotypes were used: *w^1118^* (ctrl.), *blobby*^Null^, *blobby*^GFP^, *blobby-KDRT-4xSTOP-KDRT-ALFA*, *blobby-ALFA* For tesselation sptPALM experiments: *cac-mEOS4b* vs. *cac-mEOS4b*; *blobby*^Null^ .

### Generation of transgenic flies

The following fly generated in cooperation with Well Genetics Inc. (Taipei City, Taiwan) via CRISPR/Cas9-mediated genome editing by homology-dependent repair (HDR) using guide RNAs and a dsDNA plasmid donor according to Kondo and Ueda (Kondo and Ueda, 2013): , *blobby*^Null^ and *blobby*^GFP^.

The *blobby*^Null^ allele was produced by two consecutive deletion steps, consequently introducing a 4724-bp deletion (deletion region: +13,885 nt to +18,608 from ATG of CG42795-RA) followed by a second deletion of 5,000-bp (deleted -4,997 nt to +3 nt from Stop Codon of CG42795-RC/F).

The *blobby*^GFP^ allele was produced by knock-in the EGFP tag right after S772 (based on blobby-RG isoform). The EGFP sequence is flanked by an 8 aa-linker (AGCTGTCTCTTATACACATCTGGC) upstream and a 12aa-linker (GGCGCGCCCGGGCAGATGTGTATAAGAGACAGAGGC) downstream of EGFP. The conditional blobby-KDRT-STOP-KDRT-ALFA line was created by WellGenetics Inc. by inserting the KDRT-stop-KDRT-ALFA directly after the amino acid S772 of the CG42795-PG isoform. For the analysis of presynaptic KDRT rescue, an oK6-Gal4 driver and a KD recombinase (BL# 55791) were combined with a conditional *blobby-KDRT-STOP-KDRT-ALFA*. The 3rd instar larvae with the correct genotype were collected and dissected.

### Generation of Blobby specific antibodies

Blobby^C-Term^ specific antibody. The polyclonal antibody was raised in rabbits. Immunization of the animals was performed using a His -tagged fusion protein. The coding sequence corresponding to aa 2090 - 2422 of Blobby (CG42795, based on isoform G) was MTEKKETIKDSSSKELPEKMVINSTDVGPMDPNGKTVVLLMDNEHRASKVRRLTRANTEELEDLFQALEKQLNDRNLVKSEDGRLIRVDPKPSAEQVEQTQAISDLTKEIEDFTSAKPEEENPKEAAKEDKPEPEEPEDFDWGPNTVKHHLKRKTVYLPSTKELESRFRSLERQIKLLEDVEKIDVEQRLNEIERKIKLQYSLSHEKDLNKYLELCEGKGLDDDEPVPVETPTKEAEITTARDRSRSPGRKALATKSPYTSPSRKATIKTPHTSPTRKPIIKSPYTSPSRKSAKSPYTSPSRNRQRSPSPTR SPERKSKKSPYTSPARRKPHP.

The PCR was performed using the following primer:

Blobby C-term Fwd, 5′ GATCCCATGGGGCCAAACGGAAAAACAGTGGTGCTG and Blobby C-term Rev, 5 GATCGCGGCCGCTTACATTCGCTTGGCCAAGTTCTCCTTAG.

The PCR product was subcloned into pETM-11 (HIS-tag vector). The expression and purification of the target protein were performed in Escherichia coli under native conditions. Immunization and affinity purification of the AB containing serum was performed at BioGENES company (Berlin), using the same HIS-tagged fusion protein used for immunization. Antibody specificity was tested in western blot (Fig. 1C) and staining of *blobby*^Null^ mutants (not shown). Blobbyex8b specific antibody. The polyclonal antibody was raised in rabbits. Immunization of the animals was performed using a His-MBP-tagged fusion protein. The coding sequence corresponding to amino acids 1019-1487 of Blobby (CG42795, based on isoform G) is EIEERYQALERRISQDQPSGDRQAKYIPSTAALEERFNTLEKQLSAEKQRKELSEMEAEYPIKSERIPSTADLESRFNSLTKQMSSSESSSKTPIDLKDEDRPSGSSSKNQKDSEKTSKLHKSEEPESNTKETTGETEASDSNDSKIGEKETEQPRIKKLPSTAELEDRFNALERKMSVQKSSPSKNKKEPPDEEESKSTKEPEEPEESEKANEKTSGRQTPIAKKDSKDSDQKKSETKENQSPTKNQDEKVKVKSPKSEEMIEKETSSNPKEDSHESEAATNKKVEGNRELSSEKGDHKIKEKSEEAPGKAGKETAETKNANVKDSSKKGDSQKNEAAKTSVSQTESDLKPSSKENSTSKDAEQEKTPRKSPPSTEELEKRFNALEKQMSTTNLETTKEPDQTKPATKSQSTSAEVKTQKSMKSFDDKIKEVNVAIEKEQSRVEVEVNAEKKRKNVEEAPKNKEGDSQ.

The insert was amplified by PCR. The forward primer sequence is 5′ CCGCCATGGAAATCGAGGAACGGTATCAG 3’ and the reverse primer sequence is 5’ AAGCGGCCGCCTACTGAGAATCCCCTTCTTTGTTTTTTGGGG 3’.

The PCR product was subcloned to pETM-42 vector containing an N-terminal His-MBP tag. The expression and purification of the target protein were performed in Escherichia coli under native conditions. Immunisation and affinity purification of the AB containing serum was performed at BioGENES company (Berlin), using the same His-MBP-tagged fusion protein used for immunization. Antibody specificity was tested in western blot (S 1C) and staining of *blobby*^Null^ mutants (not shown).

### Generation of the Unc13A specific antibody

The polyclonal antibody was raised in guinea pigs. Immunization of the animals was performed using a His -tagged fusion protein. The coding sequence corresponding to aa 1337 - 1632 of Unc13A was: NILPIGPQATGKKLPTVNGKSALLIKQMPTEVYDDESDTDELDVSPSTGKVPSYSIYSEQEDYYMDLQQTTPSIQPNGFYEQVNNGYDYREDYFNEEDEYKYLEQQREQEEHNQPKNKKYLKQAKISKIQPPSLDFIDVGQDDDFIYDNYHSEDDSGNYLEGSSSGSVGPIEGSIIKVDSNIEASFASLNKKSDSFTPTNDSLQKHDTVIGESTTKLTRLRTEKMCPDVDEEDENLSDHVSDLTDLSKLISQKKKTLLRGETEEVVGGHMQVLRQTEITARQRWHWAYNKIIMQLN.

The PCR was performed using the following primer:

Unc13A_AK5 Fwd, 5′ GATCCCATGGGGAACATACTCCCGATTGGCCCGCAGGC and Unc13A_AK5 Rev, 5 GATCGCGGCCGCTTAATTAAGCTGCATGATTATTTTATTG.

The PCR product was subcloned to pETM-11 (HIS-tag vector). The expression and purification of the target protein were performed in Escherichia coli under native conditions. The HIS-tagged fusion protein was used for the affinity purification of the AB-containing serum, which was obtained from Selbaq. 19GP02 Antibody specificity was tested in western blot and staining of in Unc13A mutants (data not shown).

### Western Blot analysis of adult brain protein extract

Western Blot analysis were performed as previously reported 4with some modifications. In brief, brains were dissected in HL3, homogenized in lysis buffer (0.5% Triton X-100, 2% SDS, 1× protease inhibitor, 1× sample buffer in PBS) followed by full-speed centrifugation at 18°C. One braińs supernatant was loaded to SDS-PAGE and immunoblotted according to standard protocols. The following primary antibodies were used: rabbit anti-Blobby^C-Term^ (1:500), rabbit anti-Blobbyex8b (1:500),mouse anti-Tubulin (Sigma T9026, 1:100,000).

### Isolation and purification of *Drosophila* synaptosomes

The procedure involves decapitation of adult flies (sieving, ∼6,000 heads), pulverization, homogenization (320 mM sucrose, 4 mM HEPES, protease inhibitors (complete, 11873580001, Roche)) and differential centrifugation (from low speed to higher speed: 1,000– 15,000 g) of fly heads, which allows subsequent isolation and enrichment of presynaptic and postsynaptic components5.

### BRP Co-immunoprecipitation

BRP Co-immunoprecipitation was performed according to standard protocol as previously reported ^1, 2^. Immunoprecipitation of four replicates of each, anti-rabbit IgG control and anti-BRP-IP, were performed in parallel under same condition. Both, IgG and BRP-co-IP were eluted in (5x) sample Laemmli buffer and subsequently subjected for mass-spectrometry analysis.

### Label-free global proteomics mass-spectrometry

Label-free proteomics was performed according to standard protocols as described in 7. In brief, BRP-IP eluate (4 biological replicates) and IgG eluate (4 biological replicates) were reduced (5 mM DTT at 37°C for 60 min), alkylated (40 mM CAA at RT for 30 min), loaded on NuPAGE 4–12 % Bis-Tris SDS-PAGE and subjected to an in-gel trypsin digestion procedure (ratio of 1:20 (w/w) at 37°C overnight). The LC/MS analysis was performed using a Thermo Scientific Dionex UltiMate 3000 system connected to a PepMap C-18 trap-column (0.075 × 50 mm, 3 μm particle size, 100 Å pore size; Thermo Fisher Scientific) and an in-house-packed C18 column (column material: Poroshell 120 EC-C18, 2.7 μm; Agilent Technologies) at 300 nL/min flow rate and 180 min gradient lengths. The MS1 scans were performed in the orbitrap using 120,000 resolution. The MS2 scans were acquired in the ion trap with standard AGC target settings, an intensity threshold of 1e4 and maximum injection time of 40 ms. A 1 s cycle time was set between master scans.

### Label-free global proteomics data analysis

Raw data was searched using MaxQuant version 1.6.2.6 8 using standard settings. The number of missed cleavages allowed was set to 2, label-free quantification was enabled, and the match-between-runs option was disabled. The search was performed by target-decoy competition using the UniprotKB database of Drosophila melanogaster downloaded on May 2020 containing 42,678 entries. Using the Perseus software 9 LFQ values were log2 transformed to achieve normal data distribution. Proteins identified in at least three (out of four) replicates were considered for statistical analysis all other proteins that were detected and quantified in only one replicate were excluded. Missing data were imputed by values from a normal distribution (width 0.3 standard deviations; down shift 1.8). For statistical protein enrichment analysis in the BRP-IP, a two-sided t-test between BRP-IP and negative IgG control was used with a S0 constant of 0.1 and a permutation-based FDR of 0.05. Presented fold changes have been calculated as difference from mean values of log2 transformed intensities from BRP-IP and IgG control. Microsoft Excel was used to create Volcano plot from quadruplicates of coprecipitated protein levels from the BRP-IP compared with the IgG control. The x-axis represents the log2 fold-change value, indicating the magnitude of change, and the y axis is –log10 of the p-value showing statistical significance.

### Immunostaining

Larval filets were dissected and stained as previously reported ^3^. Briefly, third instar larvae were dissected in HL3 and fixed in either 4% paraformaldehyde in PBS (pH 7.2) for 10 min for all antibodies or in ice-cold methanol for 5 min for Unc13A antibody. Afterward, the filets were washed in PBS containing 0.05% Triton X-100 (PBST) and blocked for 1h in 10% ROTI Block (Carl Roth). The primary antibody incubation was performed at 4°C overnight. Secondary antibody incubation was carried out for 3hrs at room temperature. Immunocytochemistry was equal for both conventional confocal and STED microscopy. The following primary antibodies were used: mouse anti-BruchpilotNc82/ BRP Cterm (1:100, DSHB, catalog #nc82; RRID:AB_2314866), rabbit-anti BlobbyC-term (1:300, this manuscript), anti-rabbit Blobbyex8b (1:500, this manuscript); guinea pig-anti Unc13A (1:300, this manuscript); rabbit-anti GluRIID (1:500, Qin et al., 2005). The secondary antibodies for standard immunostaining were used at the following concentrations: goat anti-HRP-Cy5 (1:250, Jackson ImmunoResearch); goat anti-rabbit-Cy3 (1:500, Jackson ImmunoResearch 111-165-144); goat anti-mouse or anti-rabbit Cy3 (1:500, abcam, ab97035/ ab6939); goat anti-mouse or anti-guinea pig or anti-rabbit Alexa Fluor 488 (1:500, Life Technologies A11001/A11073/A11008). Larvae were mounted in Vectashield (Vector Labs). Secondary antibodies for gSTED microscopy were used in the following concentrations: Alexa Fluor594-coupled goat anti-rabbit (Invitrogen A32754, 1:300), STARRED FluoTag X2-coupled goat anti-mouse (Abberior STRED-1001-500UG, 1:300); anti-GFP STARRED Fluo Tag X4 (1:300 for STED, NanoTag N0304-AbRED-L); goat anti-mouse ATTO490LS (1:50 for STED, Hypermol Cat.#:2109-1MG).

### Confocal and STED microscopy

Time-gSTED and corresponding confocal laser scanning microscopy were performed using an Abberior Instruments Expert Line STED setup equipped with an inverted IX83 microscope (Olympus), two pulsed STED lasers for depletion at 775 nm (0.98 ns pulse duration, up to 80MHz repetition rate) and at 595 nm (0.52 ns pulse duration, 40MHz repetition rate) and pulsed excitation lasers (at 488 nm, 561 nm, and 640 nm), operated by Imspector software (16.3.15507, Abberior Instruments, Germany). The dyes STARRED, Alexa Fluor594, and ATTO490 LS were depleted with a pulsed STED laser at 775 nm. Time gating was set at 750 ps. Fluorescence signals were detected sequentially by avalanche photodiode detectors at appropriate spectral regions. 2D confocal and corresponding gSTED Images were acquired sequentially with a 100x, 1.40 NA oil-immersion objective, with a pixel dwell time of 2 μs and 10x or 30x lines accumulation, respectively, at 16 bit sampling and a field of view of 10 μm × 10 μm. Lateral pixel size was set to 20 nm. Within each experiment, samples belonging to the same experimental group were acquired with equal settings. Raw triple channel gSTED images were processed for Richardson–Lucy deconvolution using the Imspector software (16.3.15507, Abberior Instruments, Germany). The point spread function was automatically computed with a 2D Lorentz function having a full-width half-maximum of 40 nm, based on measurements with 40 nm Crimson beads. Default deconvolution settings were applied.

Confocal microscopy was performed with a Leica SP8 microscope (Leica Microsystems). Images of fixed and live samples were acquired at room temperature. Confocal imaging of NMJs was done using a z step of 0.3 μm. The following objective was used: 63× 1.4 NA oil immersion for NMJ confocal imaging. All confocal images were acquired using the LCS AF software (Leica Microsystems). Images from fixed samples were taken from third instar larval NMJs (segments A2−A4). Images for figures were processed with ImageJ software to enhance brightness using the brightness/contrast function. If necessary, images were smoothened (0.5 pixel Sigma radius) using the Gauss blur function. Confocal stacks were processed with ImageJ software (http://rsbweb.nih.gov/ij/). Quantifications of AZ spot number, density and size (scored via BRP) were performed as described in ^4, 5^.

### gSTED peak-to-peak distance analysis

Deconvolved 8-bit gSTED images were used for quantification of peak-to-peak distances by line profile measurements. Line profile measurements of distances between spots were performed in Imagej (version 1.52p, NIH). Well-defined side view synapses were manually traced with the line profile tool (thickness 9 pixels/180 nm) and peak intensities across the line were retrieved using the ImageJ Macro (Macro_plot_lineprofile_multicolor from Kees Straatman, University of Leicester, Leicester, UK). Intensity values from individual synapses were exported to Excel. Local maxima were calculated with the SciPy “argrelmax” function, as described in Brockmann ^6^, in order to obtain peak intensities for different image channels and peak-to-peak distances. Only highest maxima were selected ^7^. Values were then averaged per animal.

### Colocalization analysis

Pearson’s correlation coefficient and Manders’ coefficients were used to determine the degree of co-localization on 8-bit confocal images acquired in parallel to the STED images, using the ImageJ (version 1.52p, NIH) plugin “Coloc 2”, with the PSF set to eleven pixels (220 nm) and the Costes randomizations set to ten. The Pearson’s correlation coefficients above threshold, the Manders’ tM2 coefficients of channel 2 (Rab3) with channel 1 (BRPNc82) above auto-threshold of channel 1 were measured in a given ROI (53 × 53 pixels) selecting a single bouton within an image.

### Perimeter: Segmentation and characterization of the AZ nanostructure

AZs were segmented using a custom ImageJ script (available at https://github.com/ngimber/BruchpilotSegmentation) based on the BRP signals from gSTED images. For AZ identification, images were processed by applying a Fourier bandpass filter for medium-to-large structures (0.1-2 µm). The AZs were then identified using the built-in ‘MaxEntropy’ auto-thresholding algorithm and ’watershed’ algorithm from ImageJ ^8^. Small clusters, likely representing immature AZs, were identified by applying a Fourier bandpass filter for small structures (0-0.06 µm) to the original images and applying the ‘Minimal’ threshold algorithm ^8^ from ImageJ. These structures were excluded from the AZ quantification. The perimeters of identified AZs were measured, and the results were plotted as Mean ± SEM from 22-26 boutons (4-5 animals).

### Electrophysiology

Two-electrode voltage clamp (TEVC) recordings were performed essentially as previously reported ^2^. They comprised spontaneous recordings (miniature excitatory junction currents: mEJCs, 90 s), single evoked (evoked excitatory junction currents: eEJCs, 20 repetitions at 0.2 Hz) and high-frequency recordings (paired-pulse 10 ms or 30 ms interstimulus interval, PP10 or PP30, 10 repetitions at 0.2 Hz; 60 pulses at 100 Hz for cumulative quantal content computation). All experiments were performed on third instar larvae raised at 25°C. The dissection and recording medium were extracellular haemolymphlike solution 3 ^9^; composition in mM: 70 NaCl, 5 KCl, 20 MgCl2, 10 NaHCO3, 5 trehalose, 115 sucrose, and 5 Hepes, pH adjusted to 7.2). Dissection was performed in Ca2+-free HL3 medium at room tempreature, while mEJC, eEJC, and high-frequency recordings were performed in 1.5 mM Ca2+ HL3 at room temperature. For all physiological recordings, intracellular electrodes with a resistance of 20−35 MΩ (filled with 3M KCl) were placed at muscle 6 of the abdominal segment A2/A3. The data acquired were low-pass filtered at 1 kHz and sampled at 10 kHz. The command potential for mEJC recordings was −80 mV, and −60 mV for all other recordings. Only cells with an initial membrane potential between −50 and −70 mV and input resistances of ∼4 MΩ were used for further analysis. The eEJC and paired-pulse traces were analyzed for standard parameters (amplitude, rise time, decay, charge flow, paired-pulse [PP]-ratio) by using a semiautomatic custom written Matlab script (Mathworks, version R2009a). The 100-Hz trains were analyzed for amplitudes by using a semiautomatic custom-written Matlab script that calculates eEJC amplitudes by measuring peak to baseline directly before the onset of the response. The quantal content of each response was calculated by dividing the amplitude by the mean quantal size of the respective genotype. Release-ready vesicles (y intercept) and refilling rate (slope) were determined by back extrapolation of the last 300 ms of cumulative quantal contents. Asynchronous release is estimated as the difference of the total eEJC amplitude per stimulus measured from the pre-train baseline minus the synchronous eEJC release amplitude per stimulus (measured from before the specific eEJC onset). The mEJC recordings were analyzed with pClamp 10 software (Molecular Devices). GraphPad Prism v5.01 (GraphPad Software, Inc.) was used for all fitting procedures. Data were analyzed using GraphPad Prism v8.4.2. Data distribution is normal following the D’Agostino and Pearson omnibus normality test. If data had a normal distribution, an unpaired, two-sided t test was applied for comparison of two conditions or a one-way ANOVA using Tukey post-test if more than two conditions were compared. If data did have a normal distribution, the nonparametric, two-tailed Mann–Whitney test was applied for comparison of two conditions or the nonparametric Kruskal–Wallis test with the Dunn’s post-test if more than two conditions were compared. For standard TEVC analysis, n = 1 cell, and one or two cells from four to six animals were analyzed.

### Single-particle tracking PALM

Live sptPALM experiments were conducted on male third instar larvae of *cac*-mEOS4b and *cac-*mEOS4b; *blobby*^Null^ animals at a recording temperature was set to 25°C (Fig. 6). Larval body wall preps designated for single particle tracking were prepared according to the previous studies ^10–12^ For detailed information see ^12^. and imaging experiments were performed using an inverted total internal reflection fluorescence (TIRF) setup. The microscope (Nikon Eclipse Ti) was equipped with a 100x NA 1.49 Apo TIRF oil objective (Nikon). Up to 10.000 images were captured using an EMCCD camera (iXon+ 897, Andor Technology) controlled by NIS-Elements (Nikon) at a frame rate of 20 Hz. Larval NMJ boutons were focused on the live HRP-488 staining used as an indicator for the Z-plane. We used a 1.6 magnification lens to reduce the pixel size to 107×107 nm.

Male larvae of endogenously tagged *cac-mEOS4b* and the *cac-mEOS4B*;*blobby*^Null^ were prepared, imaged and analyzed for the cumulative distribution of Cac Channels and BRP molecules in these AZs, as previously reported by ^12^. Male mutant and control larvae were dissected in in Ca2+ and Mg2+ free HL3.1 saline (in mM: NaCl 70, KCl 5, NaHCO3 10, Sucrose 115, Trehalose 5, HEPES 5). Subsequently, the larvae were washed briefly in HL3.1 imaging buffer containing 4 mM Mg2+ and 1.5 mM Ca2+ and subjected to a 5-minute live HRP-488 stain in the imaging buffer and were imaged at the HRP Z-plane of type 1b NMJs (from segments A2-A4 on Muscle 4 or 6/7). We performed live sptPALM imaging of Cac molecule populations at NMJ Ib boutons (muscle 4, 6, and 7) and imaged for 8.5 minutes per image at an acquisition rate of 20Hz.

Prior to localization detection the movies were drift corrected and cropped to exclude movement artefacts by using the ImageJ plugins NanoJ ^13^ or Thunderstorm ^14^ Analysis of the local channel density within confined regions was performed by cluster analysis based on Voronoï tessellation constructed from localized channels using the software package SR-Tessler ^14^.

### Tessellation Analysis of BRP localizations within AZ cluster and Nanocluster distributions

SR-Tessler software and the ImageJ NanoJ-SRRF were used to process and segment each BRP localization recorded by live sptPALM of all mutant and control images. Tessellation was performed on NanoJ-SRRF drift corrected sptPALM movies. This movie was then subjected to ThunderSTORM analysis to extract all PALM localizations of BRP molecules in an image containing an NMJ with several boutons, with each bouton containing high density localizations of BRP at AZs. This extracted map of all BRP localizations was processed in SR-Tesseler software to generate a tessellated map of all BRP localizations.

#### Cac-mEOS4b imaging

A list of the following parameters was used to select and analyze the highest-density Cac localizations at AZs that reflected the best TIRF Z focus and exemplified planar Cac localization clusters from the tessellation map of all Cac channel localizations in an image automatically and robustly. By this automated filtration process we excluded the out of focus and side-view orientations of synapses and their Cac clusters.

Tessellation settings: AZ boundary settings: Voronoi object: 50-200 density factor, min Area: 2000, Min Localization: 100-200 NC boundary settings: Voronoi Nanocluster object: 2 density factor, min Area: 50, Min Localization: 20. List of AZ clusters and their nanoclusters: size, area, number of localizations, density, and number of clusters were exported and filtered to remove all AZ clusters lacking a nanocluster and were also within 30-600 nm AZ and Nanoclusters limits. In addition, we applied further limitations to Cac localization number=AZ boundary: 3000 NC boundary: 500, diameter size=AZ boundary: 600-30 nm. NC boundary: 300-30 nm, Area: AZ boundary: 50000 nm2. NC boundary: 7000 nm2 and density=AZ & NC boundary: 0.072 localizations/nm2 to define well the nanodomain of Cac channels at the presynaptic membrane of the *Drosophila* NMJ. We took into consideration the parameters set for Cac localizations from recent dSTORM and QPAINT imaging studies and our previous study ^12^.

### Dendrogram

The dendrogram is constructed using the sequence similarity of all human TBC-containing proteins in conjunction with Drosophila Blobby. It is generated by aligning the entire amino acid sequence of each protein and visualized using bioinformatics software (“Geneious Prime”).

### Statistics

Data were analyzed using Prism (Version 7 & 8, GraphPad Software). Per default Student’s T test was performed to compare the means of two groups unless the data were either non-normally distributed (as assessed by D’Agostino-Pearson omnibus normality test) or if variances were unequal (assessed by F test) in which case they were compared by a Mann– Whitney U Test. For comparison of more than two groups, one-way analysis of variance (ANOVA) tests were used, followed by a Tukey’s multiple comparison test. For immunostaining, all genotypes were prepared in one session, stained in one cup and analyzed in an unbiased manner. For electrophysiological recordings, genotypes were measured in an alternating fashion on the same day and strictly analyzed in an unbiased manner. P values and n values are given in the figure legends. Means are annotated ± s.e.m. Asterisks are used to denote significance: *P < 0.05; **P < 0.01; ***P < 0.001; n.s. (not significant), P > 0.05.

