## Supplementary Figure S1 and S2 for "Identification and characterization of a synaptic active zone assembly protein"

**The PDF file includes:**

Figs. S1 and S2
References

### SUPPLEMENTARY FIGURE 1

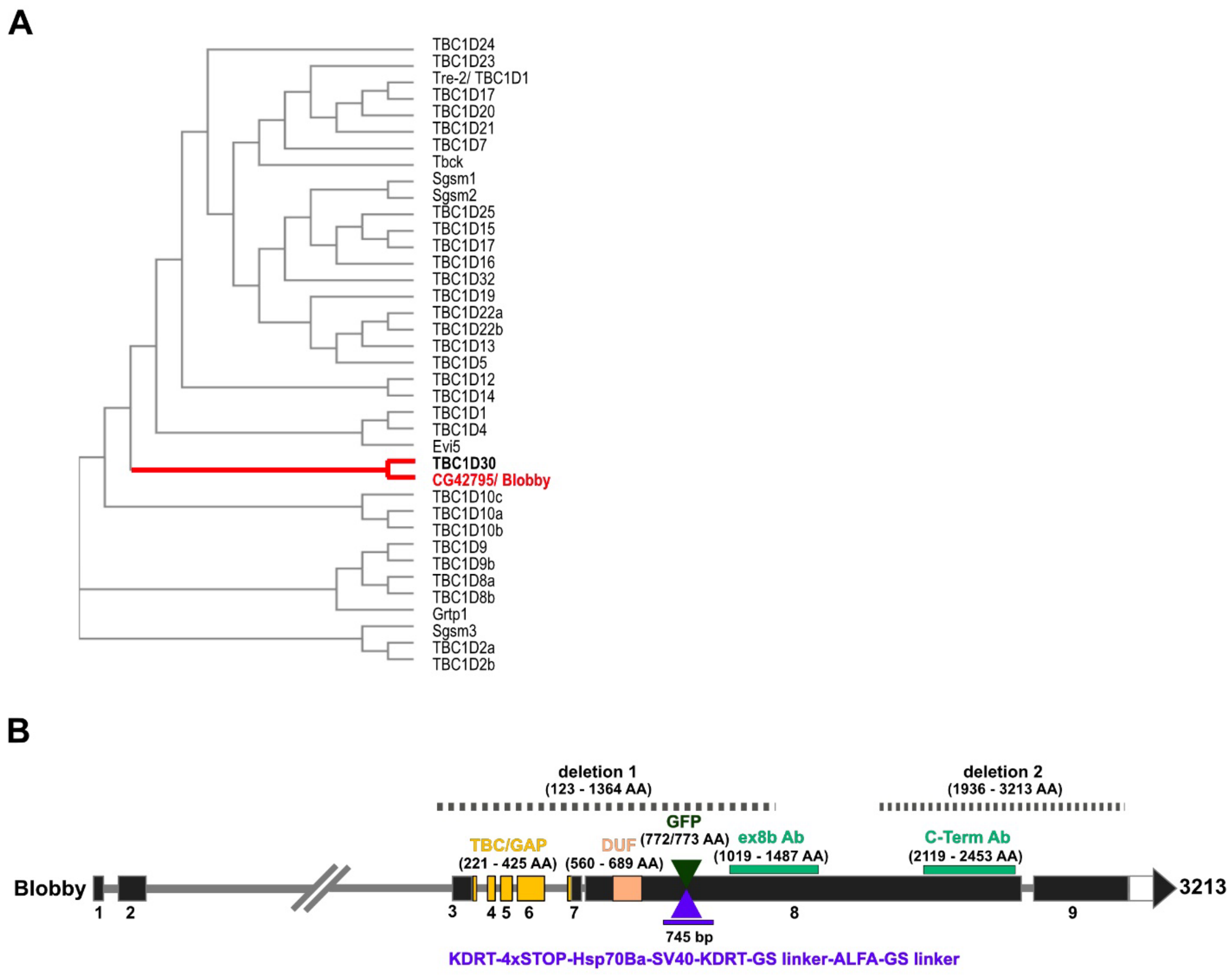


**Figure S1 | Characterization of the novel AZ scaffold protein Blobby.**

**A)** Dendrogram analysis with all predicted human TBC containing proteins in comparison with *Drosophila* Blobby. **B)** Overview map about the blobby gene locus: TBC domain (yellow), domain of unknown function (DUF, apricot), indicated antibodies (light green), position of GFP insertion (dark green), position of KDRT-STOP-KDRT-ALFA cassette insertion (purple). For generation of *blobby*$^{Null}$ two parts of the locus were deleted (dotted line; deletion 1 and 2) by using the CRISPR/Cas9 system.

### SUPPLEMENTARY FIGURE 2

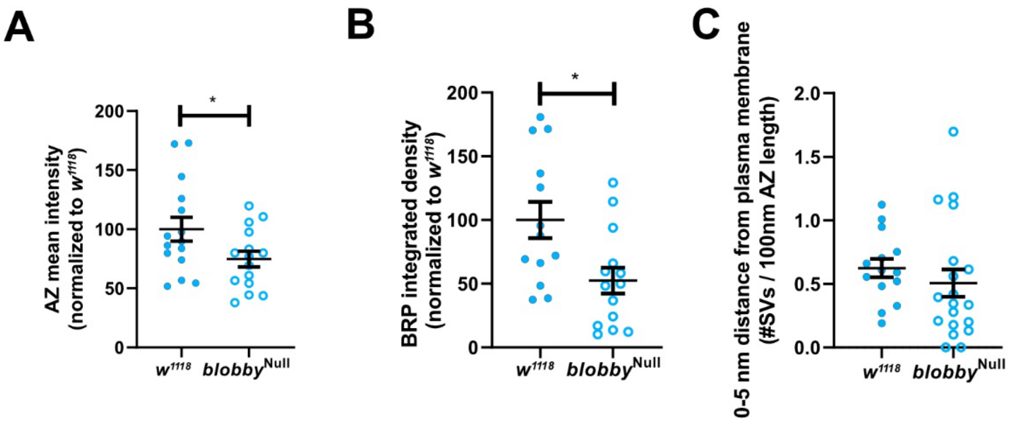


**Figure S2 | Quantification of *blobby*$^{Null}$ active zones via confocal and electron microscopy.**

**A)** Quantification of BRP$^{NC82}$ mean pixel intensity normalized to control $w^{1118}$ ($w^{1118}$ 100.00 % ± 3.80, n=32; *blobby*$^{Null}$ 72.67 % ± 3.00 n=34) and **B)** integrated density ((#BRP spots * spot area* BRP mean intensity)/ NMJ area): ($w^{1118}$ 100.00 % ± 7.12, n=25; *blobby*$^{Null}$ 67.65 % ± 7.60, n=29). Graphs in (A,B) show the mean ± SEM. n represents the number of NMJs from 5 animals per analyzed genotype. A Kolmogorov-Smirnov test was applied. *, $p < 0.05$; **, $p < 0.01$; ****, $p < 0.0001$. **C)** Quantification of synaptic vesicle density in a distance from 0-5 nm from the AZ plasma membrane per 100 nm AZ ($w^{1118}$ 0.62 ± 0.07, n=14; *blobby*$^{Null}$ 0.50 ± 0.10, n=19). Graph shows the mean ± SEM. n represents the number of boutons from 3 animals per analyzed genotype. A Kolmogorov-Smirnov test was applied.
